# Proof of concept: Malaria rapid diagnostic tests and massively parallel sequencing for surveillance of molecular markers of antimalarial resistance in Bissau, Guinea-Bissau during 2014-2017

**DOI:** 10.1101/481390

**Authors:** Sidsel Nag, Johan Ursing, Amabelia Rodrigues, Marina Crespo, Camilla Krogsgaard, Ole Lund, Frank M. Aarestrup, Michael Alifrangis, PouL-Erik Kofoed

## Abstract

Real-time and large-scale surveillance of molecular markers of antimalarial drug resistance is a potential method of resistance monitoring, to complement therapeutic efficacy studies in settings where the latter are logistically challenging. This study investigates whether routinely used malaria rapid diagnostic tests (RDTs) can be used for massive parallel amplicon sequencing. RDTs used for malaria diagnosis were routinely collected together with patient age and sex between 2014 and 2017, from two health centres in Bissau, Guinea-Bissau. A subset of positive RDTs (n=2,184) were tested for *Plasmodium* DNA content. Those containing sufficient *Plasmodium* DNA (n=1,390) were used for library preparation, consisting of amplification of gene fragments from *pfcrt, pfmdr1, pfdhfr, pfdhps* and *pfK13*. A total of 5532 gene fragments were successfully analysed on a single Illumina Miseq flow cell. Pre-screening of samples for Plasmodium DNA content proved necessary and the nested PCR protocol applied for library preparation varied notably in PCR-positivity from 13-87%. We found a high frequency of the *pfmdr1* codon 86N at 88%-97%, a significant decrease of the *pfcrt* wildtype CVMNK haplotype and elevated levels of the *pfdhfr/pfdhps* quadruple mutant ranging from 33%-51% between 2014-2017. No polymorphisms indicating artemisinin tolerance were discovered. Lastly, the demographic data indicate a large proportion of young adults (66%, interquartile range 11-28 years) presenting with *P. falciparum* infections. With some caution, our findings suggest that routine collection of RDTs could facilitate large-scale molecular surveillance of antimalarial resistance.

**Importance (word count: 147):** Continuous spread and repeated emergence of *Plasmodium falciparum* parasites resistant towards one or more antimalarials represents an enormous threat to current treatment efficacy levels, especially in sub-Saharan Africa, where 90% of malaria infections occur. In order to prevent substantial treatment failure, it is therefore recommended to monitor treatment efficacy every 2-3 years. Therapeutic efficacy studies, however, can present insurmountable logistical and financial challenges in some settings in sub-Saharan Africa. Molecular surveillance of antimalarial resistance is therefore an important proxy for treatment efficacy. However, the scale by which such studies can be performed depends on the development of high-throughput protocols and the accessibility of samples. If RDTs can be used in the high-throughput protocols available with Next Generation Sequencing (NGS)-technology, surveillance can be performed efficiently for any setting in which RDTs are already used for malaria diagnosis. The majority of settings in sub-Saharan Africa have access to RDTs.

## Background (word count (incl. conclusion): 3577)

In anticipation of novel emergence or geographic spread of especially artemisinin-resistant *Plasmodium falciparum* parasites (1-3), countries with malaria transmission are recommended to test the efficacy of their recommended artemisinin-based combination therapies (ACTs) every 2-3 years (4). Therapeutic efficacy studies are often not feasible due to economic and practical constraints in many settings in Sub-Saharan Africa (SSA). It has been suggested that molecular surveillance of genetic polymorphisms associated with antimalarial resistance could complement therapeutic efficacy studies (4-15) because these can provide early warning signs of decreasing antimalarial efficacy (5). Molecular surveillance only requires sampling of *P. falciparum* infected blood, which in turn can be acquired from finger prick samples, as used for malaria rapid diagnostic tests (RDTs) (16, 17). RDTs are now routinely used for malaria diagnostics in in SSA, and molecular analysis on used RDTs may enable large-scale molecular surveillance of antimalarial resistance (18). Large-scale surveillance also requires highly efficient and cost-effective methodologies for the genetic analysis of the parasite DNA. Novel, high-throughput protocols based on next generation sequencing (NGS) technology have the potential to achieve this (19-21).

The primary aim of this study was to evaluate collected RDTs as source of parasite DNA for NGS-based molecular surveillance of antimalarial resistance. A modified version of a recently published NGS-based amplicon sequencing methodology was used (20). The molecular approach investigated SNPs and genes associated with tolerance/resistance towards the majority of currently available antimalarial treatments in a high-throughput manner, applying massive parallel sequencing and a custom-made sample-indexing approach. A secondary aim was to provide temporal molecular marker data from the setting in Bissau, Guinea-Bissau from samples obtained from May 2014 until April 2017, as well as a basic description of the demographic trends amongst malaria patients versus non-malaria patients in the study area, by collecting minimal patient information together with the RDTs.

## Results

### Evaluation of the applicability of RDTs as source of DNA for NGS-based molecular surveillance

#### PCR-corrected RDT positivity and negativity

In total, 14,933 RDTs were used to diagnose patients at the two health centres between May 2014 and April 2017, and collected. Out of these, 2,832 RDTs were positive. A flow chart depicting the sample screening and selection process is shown in Figure 1. All positive RDTs collected at the Bandim health centre (and not at the Belem health centre) which were received in Denmark (n=2,184, 77 % of the RDT positive samples) were subjected to DNA extraction. Samples that were successfully found logged in the RDT database (n=1,879, 86 % of the DNA-extracted samples) were then checked for PCR-positivity of the ribosomal 18S *Plasmodium* subunit. The overall PCR-corrected positivity amongst these samples was 74 % (n=1,390). Median age, sex-distribution and season of collection indicated no trends in the occurrence of false positive RDTs (data not shown). A total of 304 negative RDTs from the 2014 and 2015 transmission periods were also tested for PCR-positivity. Only 1 % of these (n=3) were found to be PCR-positive.

**Figure 1.**
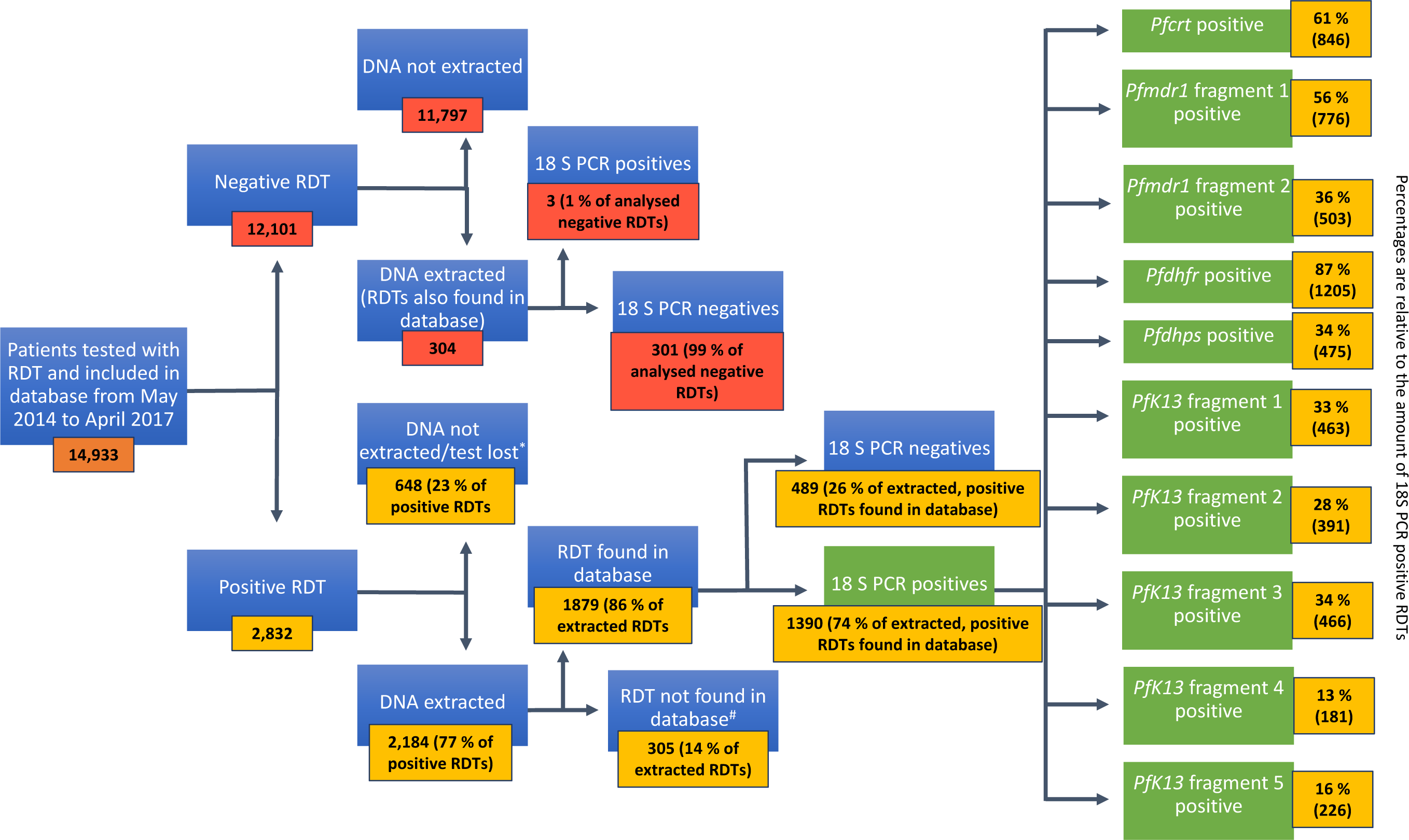
Sample screening and processing Samples were collected from all patients tested with an RDT, and subsets of positive and negative samples were used for DNA extraction and subsequent analyses. * Almost all samples from Belem were lost due to faulty extraction procedures, and other samples logged in the database were never identified amongst the RDTs received. # A number of positive RDTs received in Denmark, were not found in the database (no RDT with corresponding number or information had been logged). DNA extraction was performed before cross-referencing samples with the electronic database because the electronic database was not ready when samples were received, and postponing DNA-extraction was avoided.

#### PCR success-rate for single copy genes

In total, 1,390 PCR corrected *Plasmodium* 18S-positive samples were used in nested PCRs designed to amplify the various gene fragments analysed in this study. The success-rate of single-copy gene PCRs varied from 13% to 87% (Figure 1). Specifically, the *pfdhfr* fragment was successfully amplified for 87 % of 18S-positive RDTs, while *pfcrt* and *pfmdr1* fragment 1 were amplified for 61 % and 56 %. The *pfmdr1* fragment 2, *pfdhps* as well as *pfK13* fragments 1-3 were all amplified for 28-36 %, and lastly, *pfK13* fragments 4 and 5 were amplified for 13 % and 16 %, respectively. In total, 5532 gene fragments were successfully sequenced.

### Molecular markers of antimalarial resistance

The observed frequencies of specific haplotypes are listed in Table 1 (mixed infections were omitted from haplotype analyses), while single SNP frequencies are listed in Table 2.

**Table 1.**
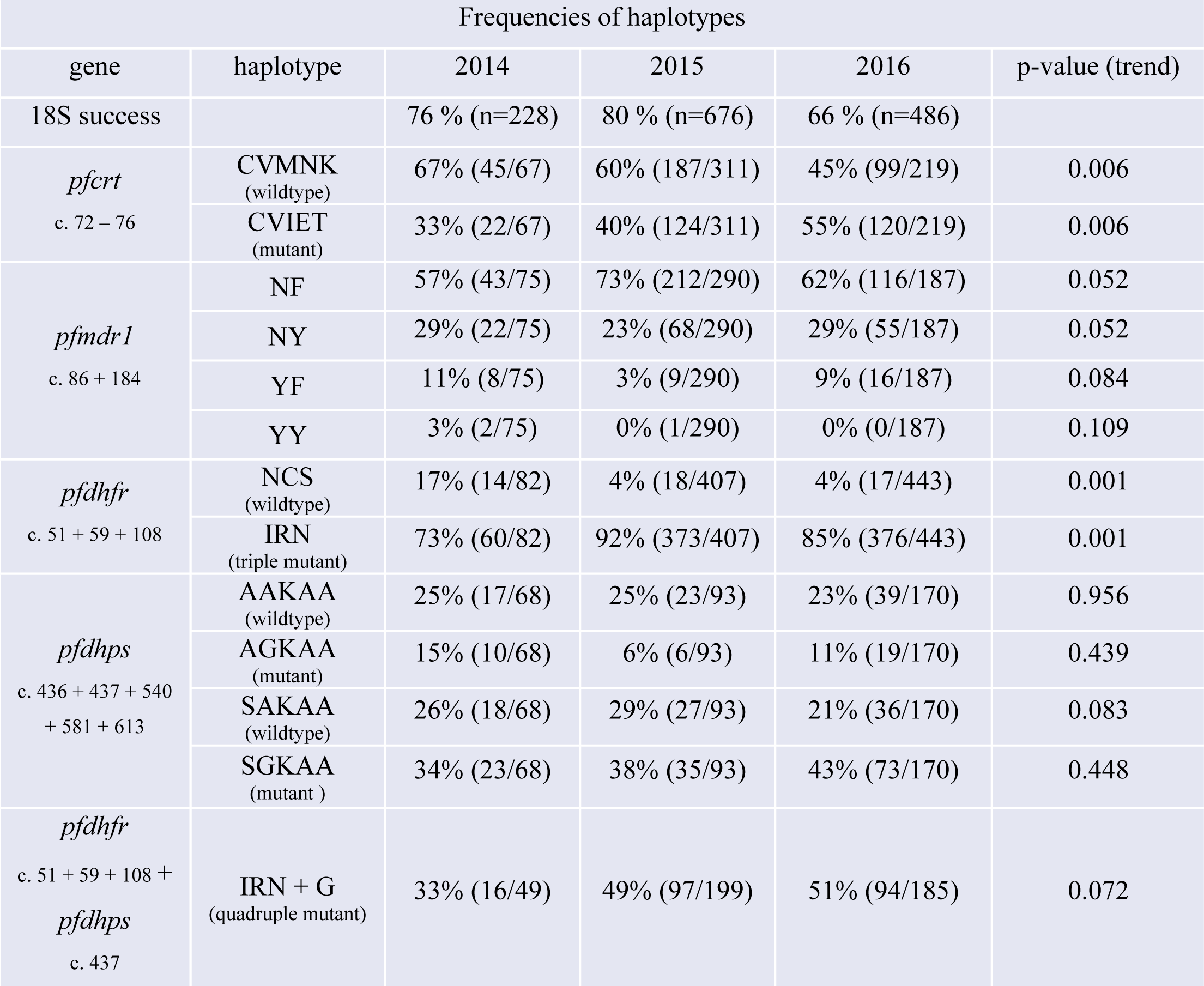
*Frequencies of haplotypes found during the transmission periods 2014-2016 (May 2014-April 2017), p-values are from Fisher’s exact test for trend over time. Mixed infections were omitted from haplotype analysis.*

**Table 2.**
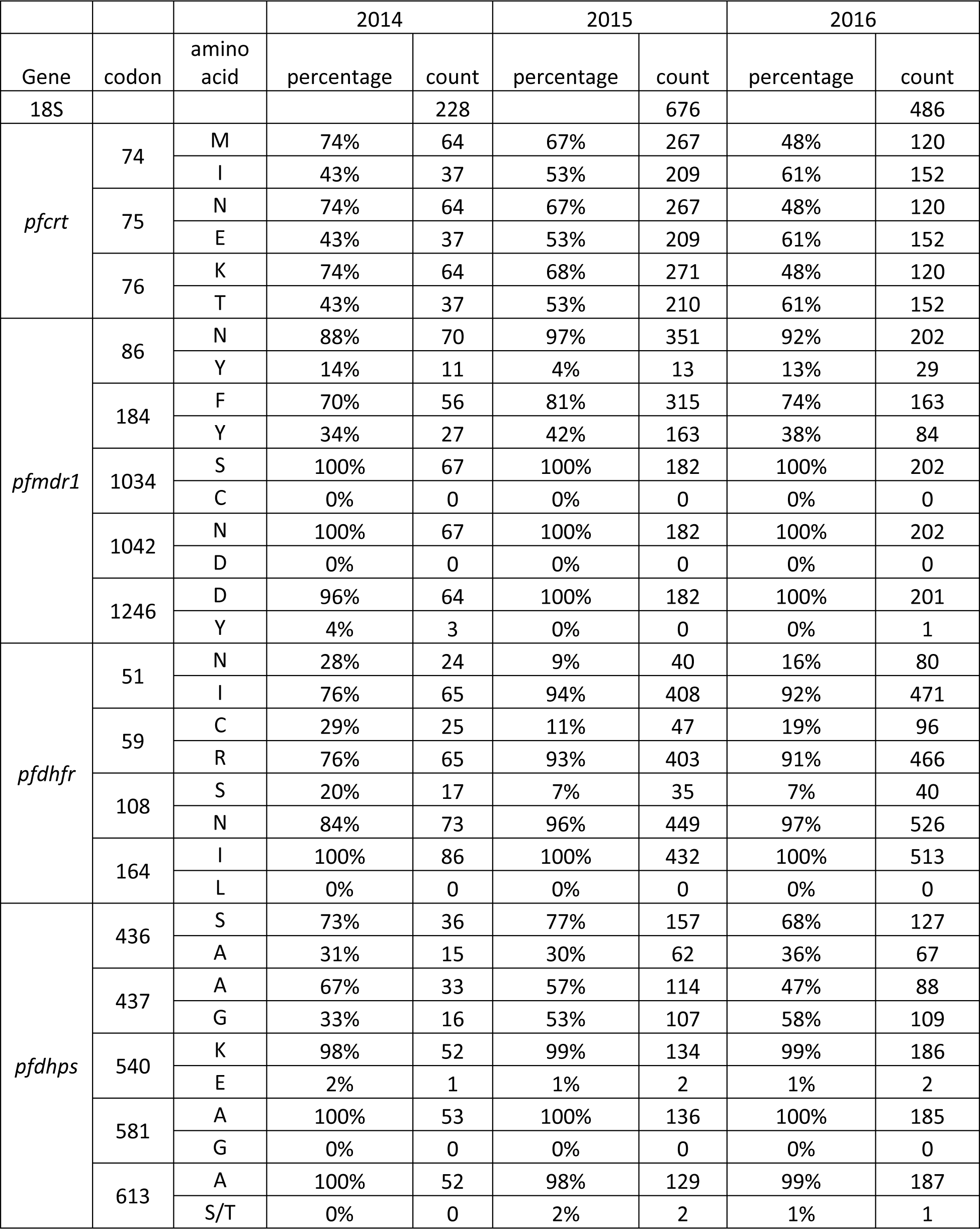
*SNP prevalence, mixed infections counted in both groups*

#### SNPs in pfcrt and pfmdr1

The *pfcrt* c. 72-76 CVMNK wild type was found to decrease significantly through the years of sampling; from 45/67 (67 %) to 187/311 (60 %) and to 99/219 (45 %) samples in 2014, 2015 and 2016, respectively (p=0.006, Fisher’s exact test) (Table 1, Figure 3A). The *pfmdr1* c. 86+184 NF haplotype was found in 43/75 (57 %), 212/290 (73 %) and 116/187 (62 %) samples, while the NY haplotype was found in 22/75 (29 %), 68/290 (23 %) and 55/187 (29 %) samples (Table 1, Figure 3B). SNP frequencies for *pfmdr1* c. 1034, 1042 and 1246 are listed in Table 2.

**Figure 3.**
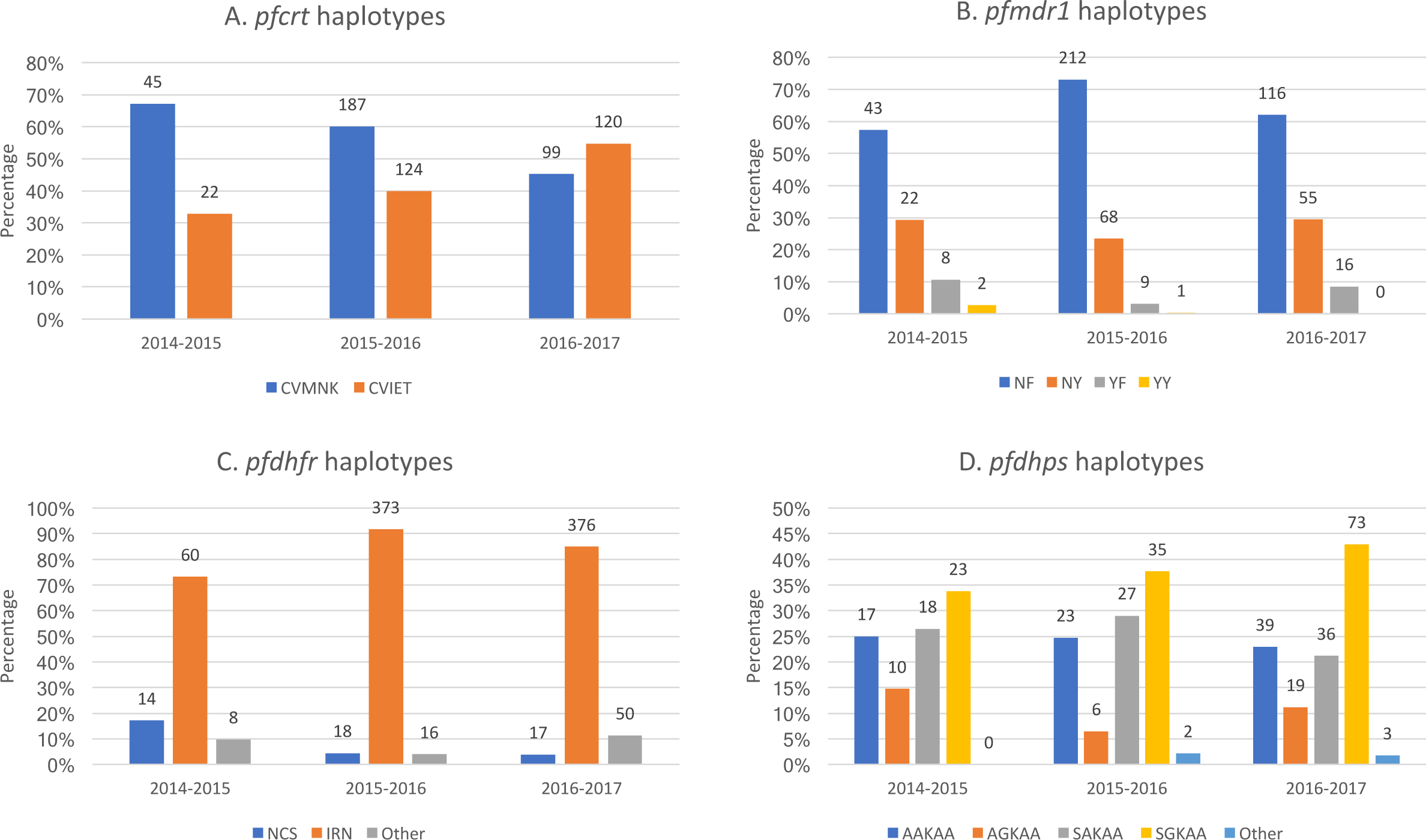
Molecular markers of antimalarial resistance 2014-2017 A) Frequency of *pfcrt* c. 72-76 haplotypes CVMNK and CVIET found each consecutive transmission season. B) Frequency of *pfmdr1* c.86 + 184 haplotypes NF, NY, YF and YY found each consecutive year. C) Frequency of *pfdhfr* c. 51 + 59 + 108 haplotypes IRN, NCS and “other” (consisting of NCN, ICN and NRN) found each consecutive year. The single mutant NCN represented 6/8 “other” *pfdhfr* haplotypes during the 2014 transmission season, while the two double mutants ICN and NRN combined accounted for 15/16 and 49/50 of “other” *pfdhfr* haplotypes found during the 2015 and 2016 transmission seasons, respectively D) Frequency of *pfdhps* c. 436 + 437 + 540 + 581 + 613 haplotypes AAKAA, AGKAA, SAKAA and SGKAA found each consecutive year.

#### SNPs in pfdhfr and pfdhps

The *pfdhfr* c. 51 + 59 + 108 IRN triple mutant was found in 60/82 (73 %), 373/407 (92 %) and 376/443 (85 %) samples in 2014, 2015 and 2016, respectively (p=0.001, Table 1, Figure 3C). The *pfdhps* c. 437G was found in 16/49 (33 %), 105/199 (53 %) and 109/185 (58 %) samples during 2014, 2015 and 2016, respectively (mixed infections included). Accordingly, the quadruple *pfdhfr/pfdhps* mutant (*pfdhfr* IRN + *pfdhps* c. 437G) was found in 33 %, 49 % and 51 % of samples. SNP frequencies for *pfdhps* c. 436, 540, 581 and 613 are listed in Table 2.

#### SNPs identified in *pfk13*

As PCR-positivity for *PfK13* fragments was very low, only data from the latest of the transmission periods is presented. A total of 311 samples collected during the 2016 transmission period were partially or completely sequenced in *pfK13*, whereof 97 were successfully sequenced in the propeller region. In total, 18 SNPs were identified in *pfK13*, only 3 of which were situated in the propeller region, 2 of which are non-synonymous (R529K and T535M) (Figure 4). In the N-terminal region, we identified 15 SNPs, 12 of which were non-synonymous (Figure 4). None of the identified SNPs occurred in more than two samples.

**Figure 4.**
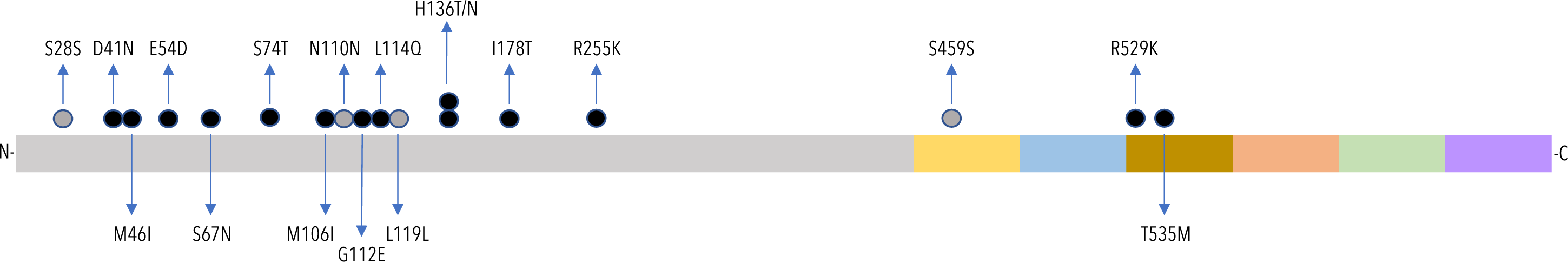
*pfK13* polymorphisms observed 2016-2017 Polymorphisms detected in *pfK13* during the transmission season from September 2016-January 2017. The grey bar indicates the N-terminal part of the translated K13 protein, while the coloured bars (yellow, blue, brown, peach, green and purple) indicate blades 1-6 in the propeller region. Grey circles indicate a synonymous SNP, while black circles indicate a non-synonymous SNP. Positions refer to amino-acid positions in the translated protein. The R529K and T535M mutations were each found only once.

### Demographic trends of RDT-positive versus RDT-negative patients

Sampling was carried out for 36 months, starting May 2014. Transmission periods were therefore defined as periods of 12 months going from May one year up to and including April the following year, which includes the high transmission period September to January. In order to compare years and transmission periods, transmission periods have been named according to the year when transmission started. The number of positive RDTs collected during the 2014, 2015 and 2016 transmission periods were 497, 1374 and 961, respectively (Figure 5A). The number of positive RDTs collected during the 2014 transmission period was substantially lower than the numbers collected in the two later transmission periods. Unexpected “dips” in the number of RDT positive patients were seen during January and September 2016 (Figure 5A).

**Figure 5.**
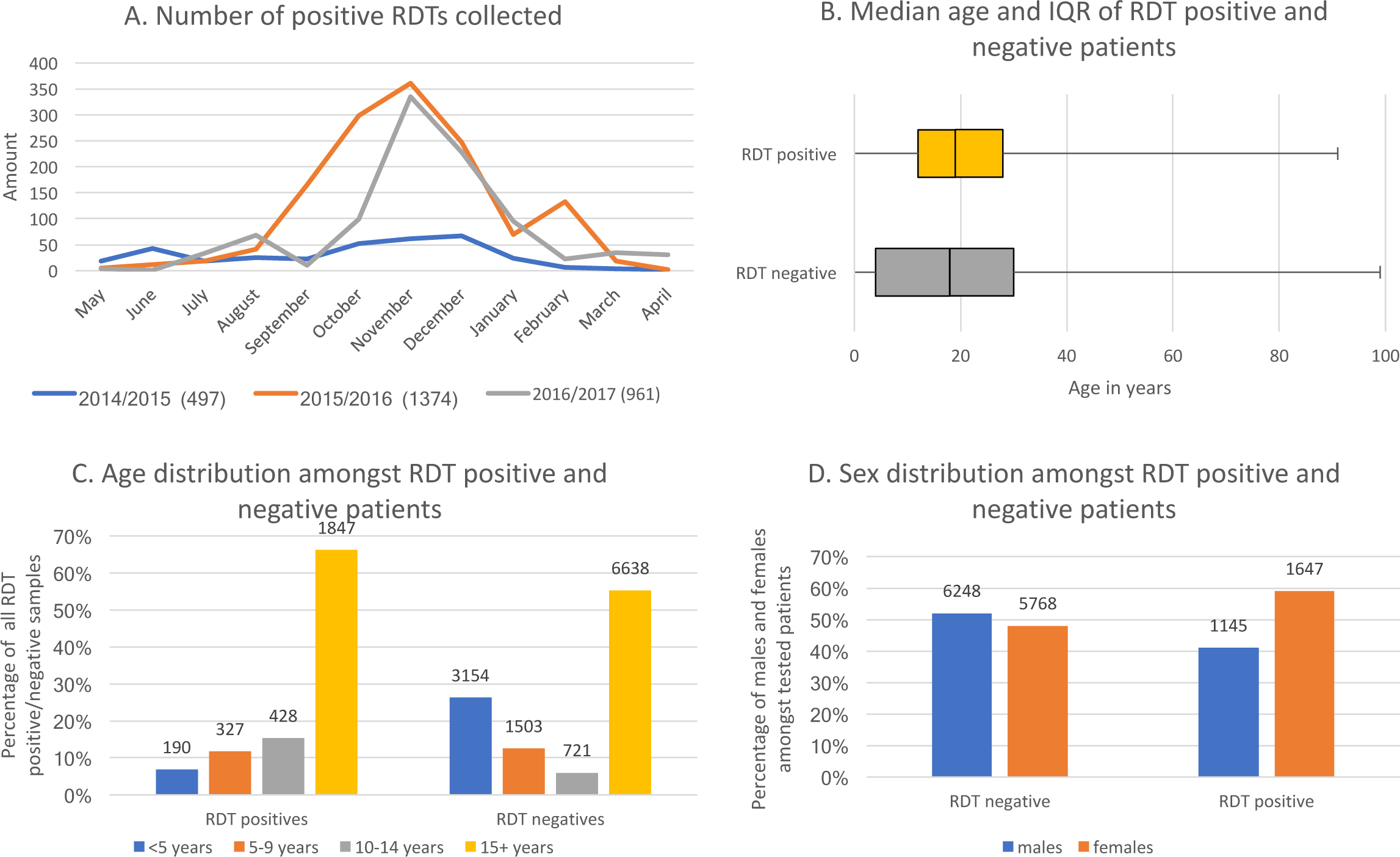
Description of RDT positive and negative patients included in the study A) Number of positive RDTs collected at the two health centres combined every month, for the three consecutive years of sampling, going from May to April. The malaria transmission season goes from September through January. B) Median age and IQR of RDT positive and RDT negative patients included throughout the study. C) Age distribution of RDT positive and RDT negative patients into groups consisting of <5 years, 5-9 years, 10-14 years and ≥15 years. D) Sex distribution of RDT positive and RDT negative patients throughout the study.

The median age of patients with a positive RDT was 19 years (interquartile range (IQR) 11-28) (Figure 5B). When divided into age groups of <5 years, 5-9, 10-14 and ≥15, the number of patients with positive RDTs were 190 (6 %), 327 (12 %), 428 (15 %) (children less than 15 years of age combined = 945 (34 %)) and 1,847 (66 %) (Figure 5C, 40 samples did not have age stated). The sex distribution amongst RDT positive patients was 1,145 males (52 %) and 1,647 females (48 %) (Figure 5D, 40 samples did not have sex stated).

The total number of negative RDTs collected was 12,101. The number of negative RDTs collected during the 2014, 2015 and 2016 transmission periods were 4,001, 4,362 and 3,738 (RDT negative database only includes until February 19^th^ 2017), respectively. The median age of patients with a negative RDT was 18 years (IQR = 4-30) (Figure 5B). When divided into age groups of <5 years, 5-9, 10-14 and ≥15, the number of patients with a negative RDT was 3,154 (26 %), 1,503 (13 %), 721 (6 %) and 6,638 (55 %) (Figure 5C, 85 samples did not have age stated). The sex distribution amongst RDT negative patients was 6,248 males (52 %) and 5,768 females (48 %) (Figure 5D, 85 samples did not have sex stated).

In order to assess whether the proportion of adults was higher in the group of RDT positive patients than in the general population, proportions were compared to that of the general population of the country, estimated in 2015 as 41.7 % children below the age of 15 vs 58.3 % adults (32). The proportion of adults within the group of RDT-positive patients was found to be significantly higher than that within the general population (Pearsons chi-square, p = 0.05), while the proportion of adults within the entire group of RDT-tested patients was not.

### Discussion

The primary aim of this study was to evaluate whether used RDTs sampled from health centres in Bissau could be applied for molecular surveillance of antimalarial resistance using a recently developed NGS protocol (20). The secondary aim was to provide temporal molecular marker data from the setting in Bissau, Guinea-Bissau from samples obtained between May 2014 and April 2017 and as well explore basic demographic trends related to malaria epidemiology in Bissau, by collecting limited patient information together with the RDTs.

### Proof of concept

Approximately 74 % of the positive RDTs were found PCR positive for the multicopy *Plasmodium* 18S subunit, indicating firstly a diagnostic false positivity percentage of 26 %, and secondly that only a maximum of 74 % of the positive RDTs collected would contain *Plasmodium* DNA, which is required for molecular surveillance. While the reasons for the high false positivity rate of the RDTs analysed in this study remain unknown (other studies suggest remainder antigens, substituted RDT buffer and non-targeted infections (33, 34)), the results indicate that the cost-efficiency of using RDTs for molecular surveillance can be affected substantially by pre-screening the samples for the presence of *Plasmodium* DNA. Furthermore, the PCR-positivity of the single-copy genes involved in conferring resistance towards antimalarial drugs varied tremendously from 13%-87% after corrected 18S PCR-positivity. Studies using erythrocyte-enhanced samples (20) or dried venous blood spots (not erythrocyte-enhanced) on filter paper (C. Schmiegelow, H. S. Hansson, S. Nag and M. Alifrangis, unpublished) subjected to the same protocol resulted in PCR positivity of at least 90 % for all fragments. Both the minute amount of parasite DNA available from an RDT, the DNA extraction protocol applied, as well as the state of the DNA in question (both at the time of extraction and at the time of running PCRs) may have contributed to the considerable variation in PCR positivity of single-copy genes. Preliminarily screening the DNA extracts for parasitaemia may give an indication of which samples can successfully produce resistance-data. While such an approach requires adding an extra qPCR step to the overall analysis, it would allow minimising reagent costs and time spent during downstream steps.

Overall, the analysis became more expensive per sample when using RDTs, than it would have been if samples had consisted of dried venous blood, not considering sampling costs. It was, however, still feasible to sequence 5532 gene fragments of approximately 500 bp, all with individual indices allowing trace-back to the sample of origin, on a single Miseq V3 flow cell with paired-end reads. Due to the possibility of simultaneously analysing a very large number of samples, the NGS protocol tested in this study has therefore proven highly affordable and also seems to remain efficient, compared to many other methods allowing trace-back to sample of origin, despite a very varied PCR-success for resistance-conferring genes. If the actual sampling costs are taken into consideration, the entire per sample cost still remains far cheaper than for dried venous blood samples, due to such samples requiring further sampling materials, labour and logistics.

Other noteworthy limitations of the current study, when considering the concept of large-scale surveillance based on routine sampling of RDTs, include the fact that routine sampling of RDTs is completely dependent on RDT availability. In our study, RDTs may have been out of stock during January and September 2016, where unexplained “dips” in malaria frequency are seen for periods of time, in which case inclusion numbers for these months would be biased. Such bias can only be assessed if logs are kept by the clinics regarding their RDT availability, along with potential use of expired batches of RDTs (which was not the case in our setting). Furthermore, the nested PCR protocol which is required for the DNA extracted from RDTs, poses a much larger contamination risk during PCR procedures, than a simplex PCR protocol (35, 36). Finally, there are no sample backups when sampling RDTs, which may become a logistical and ethical concern.

#### Molecular markers of antimalarial resistance

The high prevalence of *pfmdr1* 86N in the current study, resembles previously published data for the same study area in 2010-2012 (approximately 80 %) (37), indicating a relatively stable prevalence. The data corresponds well with the use of AL and the AL-derived selection of the *pfmdr1* c. 86 N (10, 38, 39). Importantly however, a recently performed efficacy study indicates that the efficacy of AL is still 94-95 % (25), indicating that the prevalence of the *pfmdr1* c. 86 N at levels between 88-97 % is not affecting AL treatment efficacy in this setting. AL (lumefantrine specifically) has also been shown to select for the *pfcrt* 76K wildtype (40). However, our study found a significant increase of the mutant *pfcrt* CVIET haplotype over the study period. A similar trend has previously been observed in the same study area and QN usage was speculated to be the cause (37, 41). However, it may also be that the two observed events (2010-2012 and 2014-2016) of increasing levels of the CVIET haplotype represent “highs” in a more long-term fluctuation of this haplotype.

The levels of the *pfdhfr* IRN triple mutant found in this study (fluctuating between 73-92 %) indicate selection of this haplotype since earlier studies were conducted (in 2004; prevalence of 41 %) (42). Likewise, the current levels of the *pfdhfr/pfdhps* quadruple mutant (33-51 %) indicate selection since previous studies were conducted (15 % quadruple mutant in 2004) (42). Large scale use of IPTp may have contributed to this selection, since IPTp is the only SP-based treatment that is still recommended and implemented in Guinea-Bissau (24), apart from a very recent deployment of seasonal malaria chemoprevention (SMC by use of SP+amodiaquine) in a northern region of the country (43). SP was never first-line treatment in Guinea-Bissau, but was recommended as second-line treatment from 1996-2007. Selection may also be caused by use of SP for self-treatment of malaria, the use of sulfamethoxazole-trimethoprim for bacterial infections, and finally it is also possible that quadruple mutants are imported from neighbouring countries where SP has been used as first-line treatment and where mutant haplotypes have historically been more prevalent than in Guinea-Bissau (44-46).

Importantly, the current study revealed no SNPs of concern in *pfK13* (47). Combined with the previously published data regarding *pfK13* polymorphisms from the area (20), there are no signs of artemisinin selective pressure of the kind seen in South-East Asia and Suriname (3).

### Demographic trends

The demographic data obtained from routinely collecting used RDTs, indicate that there was less malaria during the 2014 transmission period, than during 2015 and 2016 transmission periods. According to rainfall data obtained from Bandim for the three seasons, an increase in rainfall from 2014 to 2015 was observed, which may have contributed to a rise in malaria cases (yearly rainfall was 941.2 mm in 2014, 1393 mm in 2015 and 983.9 mm in 2016) (48). A country-wide long-lasting insecticide treated bed net (LLIN) distribution campaign and subsequent follow-up study carried out in June and December 2014, confirmed the low prevalence in 2014 (1.3 % amongst children aged 0-59 months and 0.7 % amongst children 5-14 years) (49). Furthermore, according to the inclusion data from our study, adults (patients ≥15 years) represent the majority of infections during transmission periods, with a significantly larger proportion of adults amongst RDT positive patients, than amongst the entire RDT-tested population and the general population of the country (32). These findings correspond well with the trend of increasing median age of malaria patients previously described for the study area (22, 23).

## Conclusion

This study provides proof of concept for the use of RDTs for molecular surveillance of antimalarial resistance through massively parallel amplicon sequencing with Illumina technology. Furthermore, the study provides evidence that there is a high frequency of the *pfmdr1* c. 86 N, that the *pfcrt* CVIET haplotype has increased significantly over the course of the study, that the *pfdhfr/pfdhps* quadruple mutant has increased substantially in frequency since 2004, and that there are no accumulating SNPs in *pfK13* as of May 2017 in Bissau, Guinea-Bissau. Lastly, the study provides evidence as to how routine sampling of used RDTs combined with minimal patient data, can provide insight regarding basic demographic trends amongst the malaria patients.

## Methods

### Study site

The current study was carried out in the capital of Guinea-Bissau. Malaria epidemiology in Guinea-Bissau has changed during the past decades and is now highly seasonal with epidemics occurring from September, peaking in November and lasting through January (22, 23). Children aged <5 years no longer account for the majority of malaria cases as the median age is gradually increasing (23). The 1^st^-line treatment for malaria is artemether-lumefantrine (AL) (24), which was recently shown to be effective in the Bissau area (25). Quinine (QN) is the 2^nd^- and 3^rd^-line treatment for malaria (24) and intermittent preventive treatment in pregnancy (IPTp) is implemented (24).

### RDT sampling

Positive and negative RDTs were collected from patients of all ages whom health workers suspected might be infected with *P. falciparum* (typically associated with presence of a fever within the last 24 hours), presenting at the Bandim or Belem health centres from May 2014 until April 2017. Patient age, sex and date of collection were written on the RDT and on a clinical records form. All information was put into a folder on a daily basis, and subsequently entered into an electronic database. RDTs were collected in a storage box containing silica gel, which was kept dark at room temperature, and stored between 3 and 9 months before shipment to Denmark, where they were stored between 0 and 9 months at room temperature before DNA extraction was performed.

### DNA extraction

DNA was extracted by the chelex method, as described previously (26), in a 96-well format with no samples in lane 12 and 4 blanks dispersed between lanes 1 and 11.

### PCR-corrected RDT positivity and negativity

A PCR amplifying the multicopy ribosomal 18S subunit of all *Plasmodium* species was performed on all positive RDTs received in Denmark, as well as on 304 negative RDTs from the high transmission seasons of 2014 and 2015. The PCR protocol has been described previously (27, 28).

### Genetic analysis

Molecular markers of antimalarial resistance were assessed by NGS-based amplicon sequencing, using a modified version of a previously published protocol (20). In brief, amplicons (amplified gene-fragments) from the infecting parasites, depicted in Figure 2, were produced by PCR and prepared for massively parallel sequencing on the Illumina Miseq sequencer through a previously published PCR-based library-preparation method (20) (based on Illumina’s own protocol for 16S metagenomics sequencing (29)). All amplicons pertaining to the same infection were barcoded with the same unique set of custom-made indices in the 5’ and 3’ ends. All barcoded amplicons were pooled prior to sequencing and sequenced in parallel. De-multiplexing of sequence data was performed based on all of the unique index-combinations given to the samples during library preparation. The original multiplex, non-nested amplification of gene-fragments were modified to simplex, nested amplification of the same or slightly modified gene-fragments (Figure 2) due to the minute amount of DNA contained in RDT extracts (20). As all PCRs were performed in simplex, certain fragments were redesigned to accommodate all of the genetic positions of interest within a single fragment (to reduce the number of PCRs), instead of two fragments (regarding *pfdhps* and 3’-*pfmdr1*). In these cases, certain areas of the fragments are not sequenced, as the paired-end 300 bp sequencing is not long enough to sequence the entire fragment. All primers are listed in supplementary Table 1.

**Figure 2.**
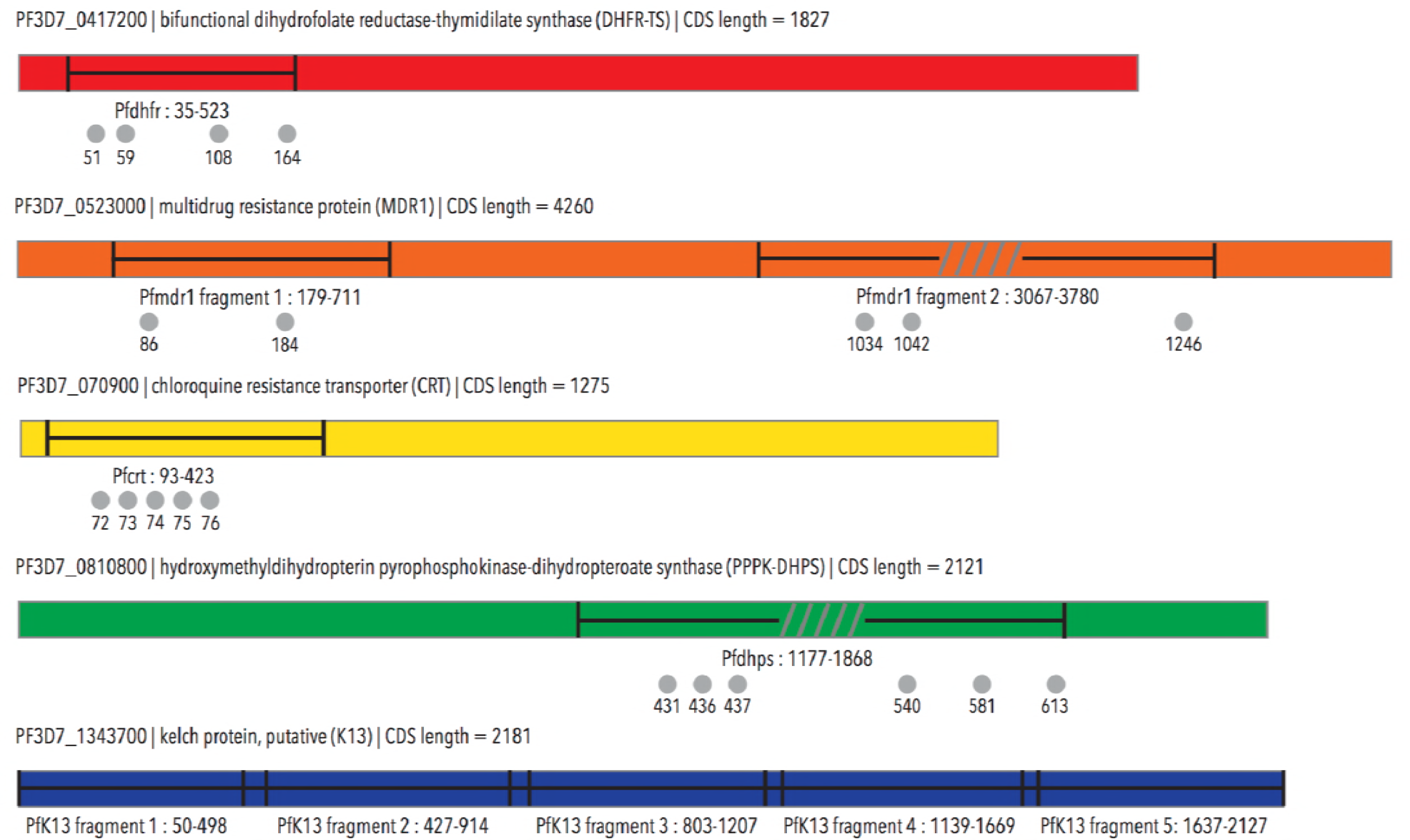
Amplicons incorporated in the sequencing library preparation Fragments from *pfcrt, pfmdr1, pfdhfr, pfdhps* and *pfK13* were incorporated in the sequencing library. Grey circles indicate SNP positions of interest, and the numbering corresponds to codons. Grey lines indicate areas of amplicons that are not sequenced.

#### Controls and duplicates

The majority of samples were run once, with 10 % of samples run as duplicates. Control samples used in the study consisted of DNA from well characterised parasites, namely 3D7, FCR3, DD2, K1, 7G8, MRA-1238 and MRA-1239 (12), the latter two of which are *pfK13* controls. Other controls consisted of patient samples from earlier studies, where the haplotypes within specific genes are known, namely AA (*pfdhps* 436A+437A), AG (*pfdhps* 436A+437G), 540E (*pfdhps* 540E) and 164L (*pfdhfr* 164L). An entire overview of control sample haplotypes is listed in supplementary Table 2.

#### Library preparation

PCRs were performed as described previously (20), with the following alterations: all fragments were amplified individually, and as nested PCRs. The outer and the nested PCR programs were identical to the previously published “gene-specific PCR”, except that they consisted of 40 cycles each. The nested PCR was performed with primers containing the overhangs, as was previously the case for the “gene-specific PCR”. The nested PCR products were pooled according to sample of origin, prior to index PCR, as described previously. The index PCR was run according to the original protocol (20). All primers and corresponding fragments are listed in supplementary Table 1, and depicted in Figure 2. Fragments included in the study cover *pfcrt* codon (c) c. 31-138, *pfmdr1* c. 60-237 and c. 1022-1260 (where c. 1123-1160 are not sequenced), *pfdhfr* c. 12-174, *pfdhps* c. 392-622 (where c. 492-522 are not sequenced) as well as *pfK13* c. 17-709.

Amplicon purification, dilution, pooling and sequencing were all performed as previously described, at the DTU Multi Assay Core (DMAC), Technical University of Denmark (20).

#### Quality trimming and base calling

Data analysis of raw sequencing reads was performed using *cutadapt* (30) and *assimpler* (31), as described previously (20). All primer sequences were trimmed from the raw data prior to SNP analysis.

### Mixed infections

Infections were defined as mixed if more than one base was called for a given position in a given sample, and was supported by at least 25 % of the base calls for that position for the sample in question.

### Statistics

Pearson’s chi-square was used to assess whether there was a difference in the proportions of children and adults amongst RDT-positive patients as compared to the general population. Fisher’s exact test was used to assess whether there was a significant trend over time in the frequencies of the detected haplotypes. Mixed infections were counted in all groups for single SNP-prevalence and omitted for haplotypes.

## Ethical approval

Ethical approval for conducting the study and sampling used RDTs was acquired from the ethical review board in Bissau (ref: 022/CNES/INASA/2014, dated September 17th 2014).

## Funding

This work was supported by Mærsk Foundation (bilagnr. OMPC1400002999), Else og Mogens Wedell-Wedellborgs Fond (J. Nr. 13-15-1) and Carl og Ellen Hertz’ Legat til Dansk Læge-og Naturvidenskab (løbenr. 6.14.2 and løbenr. 13.17.2)

### Acknowledgements

The authors would like to express great gratitude to the health workers at the Bandim and Belem health centres for collecting samples and typing patient data into the electronic database. Furthermore, the authors would like to acknowledge everybody who has travelled with extra suitcases full of RDTs from Guinea-Bissau to Denmark, as well as Ulla Abildtrup and Kenneth Kauffmann Slot for laboratory support, DMAC (DTU) and Marlene Danner Dalgaard for NGS expertise and performance and finally Frederik Shaltz-Buchholzer and Mikael Bay von Scholten for database support.

## Tables

**Supplementary Table S1.**
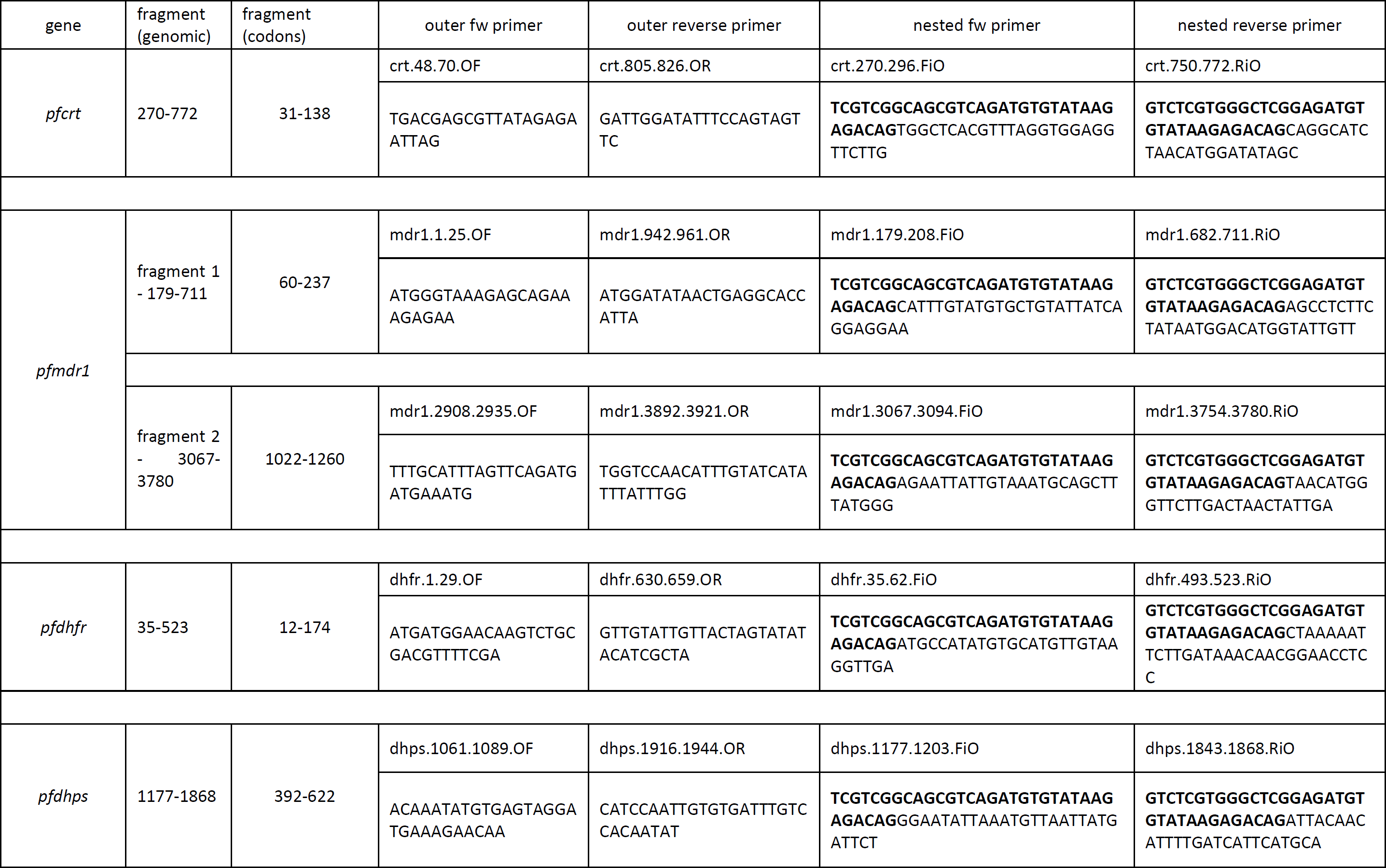
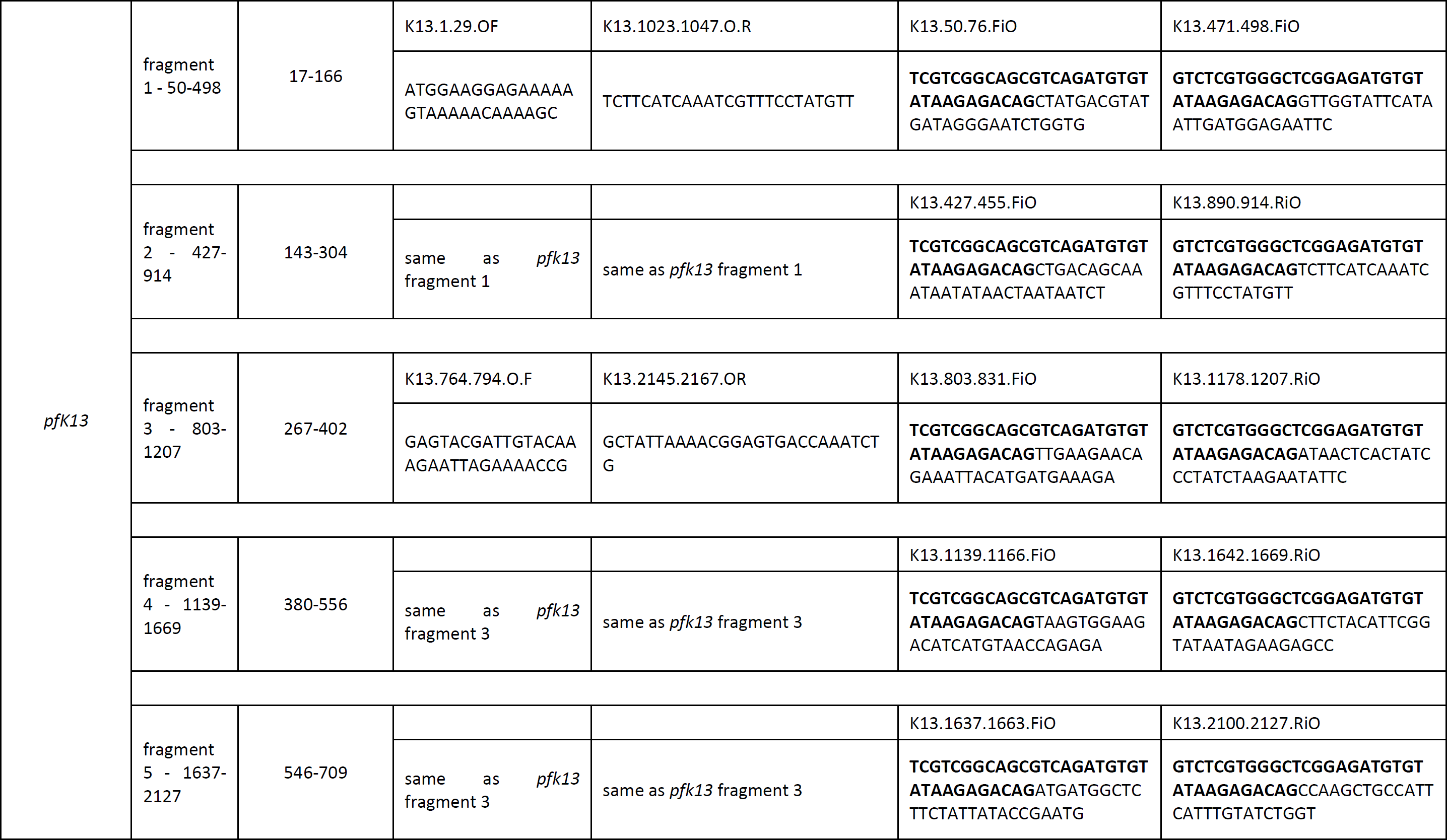
*Primers and fragments amplified*

**Supplementary Table S2.**
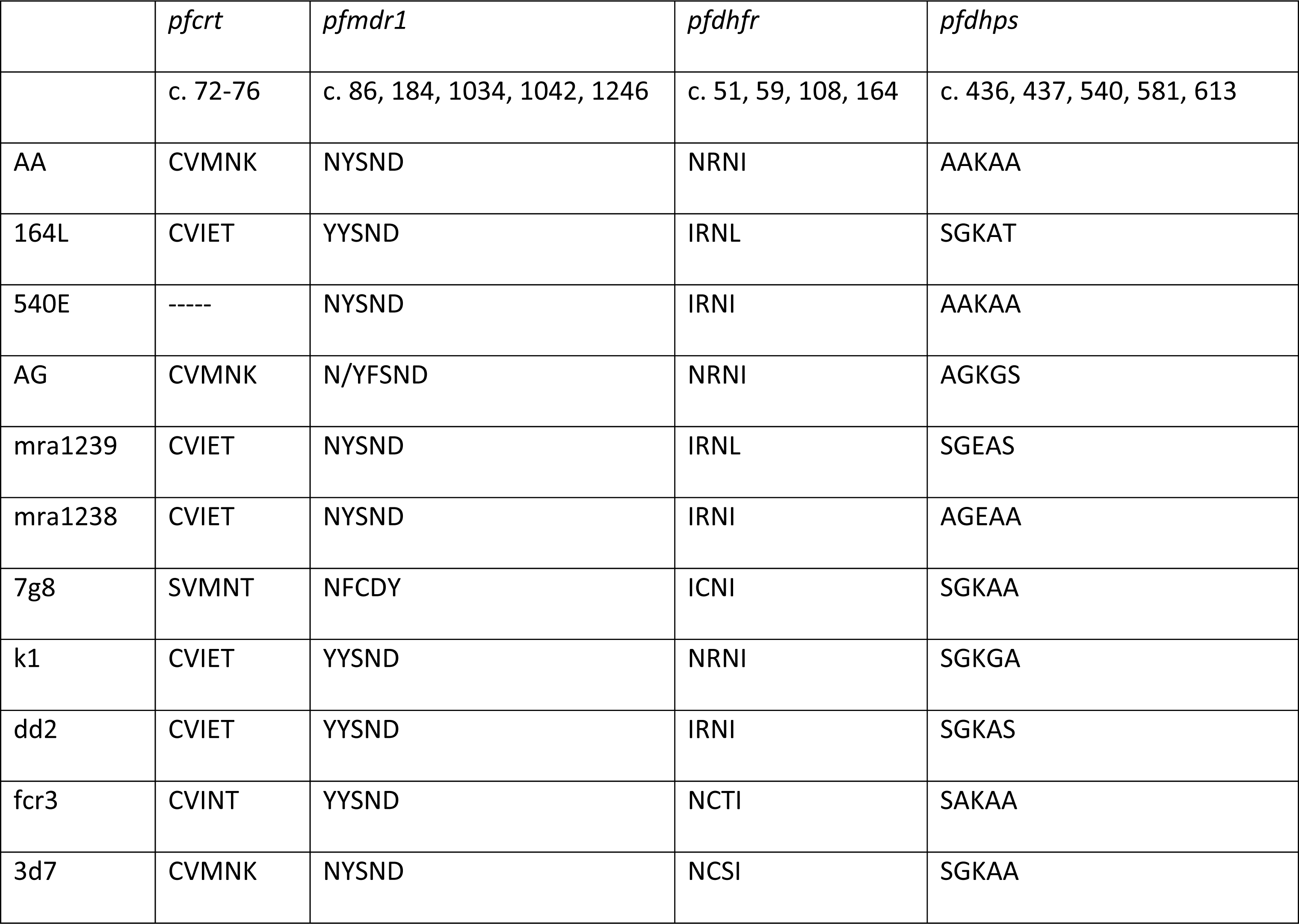
*Control sample data*

## REFERENCES

1. Muller IB, Hyde JE. Antimalarial drugs: modes of action and mechanisms of parasite resistance. Future Microbiol. 2010;5(12):1857–73.

2. Sibley CH. Understanding drug resistance in malaria parasites: basic science for public health. Mol Biochem Parasitol. 2014;195(2):107–14.

3. World Health Organization. Status Report on Artemisinin Resistance. 2014.

4. World Health Organization. Methods for surveillance of antimalarial drug efficacy. 2009.

5. Christian Nsanzabana et al. Meeting of experts on antimalarial drug resistance using molecular markers. 2018.

6. Fidock DA, Nomura T, Talley AK, Cooper RA, Dzekunov SM, Ferdig MT, et al. Mutations in the P. falciparum digestive vacuole transmembrane protein PfCRT and evidence for their role in chloroquine resistance. Mol Cell. 2000;6(4):861–71.

7. Cowman AF, Morry MJ, Biggs BA, Cross GA, Foote SJ. Amino acid changes linked to pyrimethamine resistance in the dihydrofolate reductase-thymidylate synthase gene of Plasmodium falciparum. Proceedings of the National Academy of Sciences of the United States of America. 1988;85(23):9109–13.

8. Triglia T, Menting JG, Wilson C, Cowman AF. Mutations in dihydropteroate synthase are responsible for sulfone and sulfonamide resistance in Plasmodium falciparum. Proceedings of the National Academy of Sciences of the United States of America. 1997;94(25):13944–9.

9. Price RN, Uhlemann AC, Brockman A, McGready R, Ashley E, Phaipun L, et al. Mefloquine resistance in Plasmodium falciparum and increased pfmdr1 gene copy number. Lancet. 2004;364(9432):438–47.

10. Sisowath C, Ferreira PE, Bustamante LY, Dahlstrom S, Martensson A, Bjorkman A, et al. The role of pfmdr1 in Plasmodium falciparum tolerance to artemether-lumefantrine in Africa. Trop Med Int Health. 2007;12(6):736–42.

11. Veiga MI, Dhingra SK, Henrich PP, Straimer J, Gnadig N, Uhlemann AC, et al. Globally prevalent PfMDR1 mutations modulate Plasmodium falciparum susceptibility to artemisinin-based combination therapies. Nature communications. 2016;7:11553.

12. Ariey F, Witkowski B, Amaratunga C, Beghain J, Langlois AC, Khim N, et al. A molecular marker of artemisinin-resistant Plasmodium falciparum malaria. Nature. 2014;505(7481):50–5.

13. Amato R, Lim P, Miotto O, Amaratunga C, Dek D, Pearson RD, et al. Genetic markers associated with dihydroartemisinin-piperaquine failure in Plasmodium falciparum malaria in Cambodia: a genotype-phenotype association study. Lancet Infect Dis. 2017;17(2):164–73.

14. Witkowski B, Duru V, Khim N, Ross LS, Saintpierre B, Beghain J, et al. A surrogate marker of piperaquine-resistant Plasmodium falciparum malaria: a phenotype-genotype association study. Lancet Infect Dis. 2017;17(2):174–83.

15. Christian Nsanzabana et al. Target product profile for a molecular assay for antimalarial drug resistance surveillance. 2018.

16. Ishengoma DS, Lwitiho S, Madebe RA, Nyagonde N, Persson O, Vestergaard LS, et al. Using rapid diagnostic tests as source of malaria parasite DNA for molecular analyses in the era of declining malaria prevalence. Malaria journal. 2011;10:6.

17. Morris U, Aydin-Schmidt B, Shakely D, Martensson A, Jornhagen L, Ali AS, et al. Rapid diagnostic tests for molecular surveillance of Plasmodium falciparum malaria-assessment of DNA extraction methods and field applicability. Malaria journal. 2013;12:106.

18. Ndiaye M, Sow D, Nag S, Sylla K, Tine RC, Ndiaye JL, et al. Country-Wide Surveillance of Molecular Markers of Antimalarial Drug Resistance in Senegal by Use of Positive Malaria Rapid Diagnostic Tests. Am J Trop Med Hyg. 2017;97(5):1593–6.

19. Daniels R, Ndiaye D, Wall M, McKinney J, Sene PD, Sabeti PC, et al. Rapid, field-deployable method for genotyping and discovery of single-nucleotide polymorphisms associated with drug resistance in Plasmodium falciparum. Antimicrob Agents Chemother. 2012;56(6):2976– 86.

20. Nag S, Dalgaard MD, Kofoed PE, Ursing J, Crespo M, Andersen LO, et al. High-throughput resistance profiling of Plasmodium falciparum infections based on custom dual indexing and Illumina next generation sequencing-technology. Scientific reports. 2017;7(1):2398.

21. Levitt B, Obala A, Langdon S, Corcoran D, O’Meara WP, Taylor SM. Overlap Extension Barcoding for the Next Generation Sequencing and Genotyping of Plasmodium falciparum in Individual Patients in Western Kenya. Scientific reports. 2017;7:41108.

22. Rodrigues A, Schellenberg JA, Kofoed PE, Aaby P, Greenwood B. Changing pattern of malaria in Bissau, Guinea Bissau. Trop Med Int Health. 2008;13(3):410–7.

23. Ursing J, Rombo L, Rodrigues A, Aaby P, Kofoed PE. Malaria transmission in Bissau, Guinea-Bissau between 1995 and 2012: malaria resurgence did not negatively affect mortality. PloS one. 2014;9(7):e101167.

24. World Health Organization. World Malaria Report 2016. Geneva: World Health Organization; 2017. Report No.: 978-92-4-151171-1 Contract No.: August 14th.

25. Ursing J, Rombo L, Rodrigues A, Kofoed PE. Artemether-Lumefantrine versus Dihydroartemisinin-Piperaquine for Treatment of Uncomplicated Plasmodium falciparum Malaria in Children Aged Less than 15 Years in Guinea-Bissau - An Open-Label Non-Inferiority Randomised Clinical Trial. PloS one. 2016;11(9):e0161495.

26. Schriefer ME, Sacci JB, Jr., Wirtz RA, Azad AF. Detection of polymerase chain reaction-amplified malarial DNA in infected blood and individual mosquitoes. Exp Parasitol. 1991;73(3):311–6.

27. Snounou G, Viriyakosol S, Jarra W, Thaithong S, Brown KN. Identification of the four human malaria parasite species in field samples by the polymerase chain reaction and detection of a high prevalence of mixed infections. Mol Biochem Parasitol. 1993;58(2):283–92.

28. Toure M, Petersen PT, Bathily SN, Sanogo D, Wang CW, Schioler KL, et al. Molecular Evidence of Malaria and Zoonotic Diseases Among Rapid Diagnostic Test-Negative Febrile Patients in Low-Transmission Season, Mali. Am J Trop Med Hyg. 2017;96(2):335–7.

29. Illumina. 16S Metagenomic Sequencing Library Preparation - Preparing 16S Ribosomal RNAGene Amplicons for the Illumina MiSeq System 2013 [Available from: http://www.illumina.com/content/dam/illumina- support/documents/documentation/chemistry_documentation/16s/16s-metagenomic-library-prep-guide-15044223-b.pdf.

30. Martin M. Cutadapt Removes Adapter Sequences From High-Throughput Sequencing Reads. EMBnetjournal. 2011;17(1):10–2.

31. Leekitcharoenphon P, Nielsen EM, Kaas RS, Lund O, Aarestrup FM. Evaluation of whole genome sequencing for outbreak detection of Salmonella enterica. PloS one. 2014;9(2):e87991.

32. Department of Economics and Social Affairs UN. World Population Prospects The 2017 Revision. 2018.

33. Mouatcho JC, Goldring JP. Malaria rapid diagnostic tests: challenges and prospects. J Med Microbiol. 2013;62(Pt 10):1491–505.

34. Gillet P, Mori M, Van den Ende J, Jacobs J. Buffer substitution in malaria rapid diagnostic tests causes false-positive results. Malaria journal. 2010;9:215.

35. Murray DC, Coghlan ML, Bunce M. From benchtop to desktop: important considerations when designing amplicon sequencing workflows. PloS one. 2015;10(4):e0124671.

36. Seitz V, Schaper S, Droge A, Lenze D, Hummel M, Hennig S. A new method to prevent carry-over contaminations in two-step PCR NGS library preparations. Nucleic acids research. 2015;43(20):e135.

37. Jovel IT, Kofoed PE, Rombo L, Rodrigues A, Ursing J. Temporal and seasonal changes of genetic polymorphisms associated with altered drug susceptibility to chloroquine, lumefantrine, and quinine in Guinea-Bissau between 2003 and 2012. Antimicrob Agents Chemother. 2015;59(2):872– 9.

38. Malmberg M, Ferreira PE, Tarning J, Ursing J, Ngasala B, Bjorkman A, et al. Plasmodium falciparum drug resistance phenotype as assessed by patient antimalarial drug levels and its association with pfmdr1 polymorphisms. The Journal of infectious diseases. 2013;207(5):842–7.

39. Thomsen TT, Madsen LB, Hansson HH, Tomas EV, Charlwood D, Bygbjerg IC, et al. Rapid selection of Plasmodium falciparum chloroquine resistance transporter gene and multidrug resistance gene-1 haplotypes associated with past chloroquine and present artemether-lumefantrine use in Inhambane District, southern Mozambique. Am J Trop Med Hyg. 2013;88(3):536–41.

40. Sisowath C, Petersen I, Veiga MI, Martensson A, Premji Z, Bjorkman A, et al. In vivo selection of Plasmodium falciparum parasites carrying the chloroquine-susceptible pfcrt K76 allele after treatment with artemether-lumefantrine in Africa. The Journal of infectious diseases. 2009;199(5):750–7.

41. Ursing J, Kofoed PE, Rodrigues A, Rombo L. No seasonal accumulation of resistant P. falciparum when high-dose chloroquine is used. PloS one. 2009;4(8):e6866.

42. Kofoed PE, Alfrangis M, Poulsen A, Rodrigues A, Gjedde SB, Ronn A, et al. Genetic markers of resistance to pyrimethamine and sulfonamides in Plasmodium falciparum parasites compared with the resistance patterns in isolates of Escherichia coli from the same children in Guinea-Bissau. Trop Med Int Health. 2004;9(1):171–7.

43. Médicins sans frontières. Guinea-Bissau: Many children would be saved if they arrived earlier 2016 [Available from: http://www.msf.org/en/article/guinea-bissau-%E2%80%9Cmany- children-would-be-saved-if-they-arrived-hospital-earlier%E2%80%9D.

44. Papa Mze N, Ndiaye YD, Diedhiou CK, Rahamatou S, Dieye B, Daniels RF, et al. RDTs as a source of DNA to study Plasmodium falciparum drug resistance in isolates from Senegal and the Comoros Islands. Malaria journal. 2015;14:373.

45. Ndiaye YD, Diedhiou CK, Bei AK, Dieye B, Mbaye A, Mze NP, et al. High resolution melting: a useful field-deployable method to measure dhfr and dhps drug resistance in both highly and lowly endemic Plasmodium populations. Malaria journal. 2017;16(1):153.

46. Nwakanma DC, Duffy CW, Amambua-Ngwa A, Oriero EC, Bojang KA, Pinder M, et al. Changes in malaria parasite drug resistance in an endemic population over a 25-year period with resulting genomic evidence of selection. The Journal of infectious diseases. 2014;209(7):1126–35.

47. Menard D, Khim N, Beghain J, Adegnika AA, Shafiul-Alam M, Amodu O, et al. A Worldwide Map of Plasmodium falciparum K13-Propeller Polymorphisms. N Engl J Med. 2016;374(25):2453–64.

48. World Weather Online. Weather in Bandim, Guinea-Bissau 2018 [Available from: https://www.worldweatheronline.com/lang/en-au/bandim-weather-averages/bissau/gw.aspx.

49. The National Institute of Public Health Guinea-Bissau. Evaluation of the long lasting insecticide treated net distribution campaign impact in Guinea-Bissau. 2015.

